# Individuals with clinically relevant autistic traits tend to have an eye for detail

**DOI:** 10.1101/367532

**Authors:** Arjen Alink, Ian Charest

## Abstract

Individuals with an autism spectrum disorder (ASD) diagnosis are often described as having an ‘eye for detail’^1^. This observation, and the finding that individuals with ASD tend to ‘see the trees before the forest’ when performing the Navon task^2^, has led to the proposal that ASD is characterized by a bias towards processing local image details^3^. However, it remains to be shown that natural image recognition in individuals with autism depends more on fine image detail. Here, we resolve this issue by showing that natural image recognition relies more on details in individuals with an above-median number of autistic traits. Furthermore, we found that reliance on details was best predicted by the presence of the most clinically relevant autistic traits. Therefore, our findings raise the possibility that a wide range of real-life abilities and difficulties associated with ASD are related to an enhanced reliance on visual details.

---

Our findings resulted from the application of a new experimental paradigm with which we are able to measure the relative contribution of low-level image features to image recognition, using a technique similar to reverse correlation^4,5^. During the experiment, 52 participants were presented with partial reconstructions of five cat and five dog images. To create these stimuli, we first selected 1000 Gabor wavelets (with varying position, spatial frequency and orientation) which sum provided a good estimate of pixel intensity values of the original cat and dog images. Partial reconstructions contained a random selection of 90 of the 1000 features (Figure 1). Via button presses, participants indicated whether they recognized a dog, a cat or whether they were not sure. Participants were assigned to the high AQ group (n = 25) if they had an autistic-spectrum quotient^6^ (AQ) higher than the median across all participants (AQ > 14) while the others were assigned to the low AQ group (n = 27).

**Figure 1.**
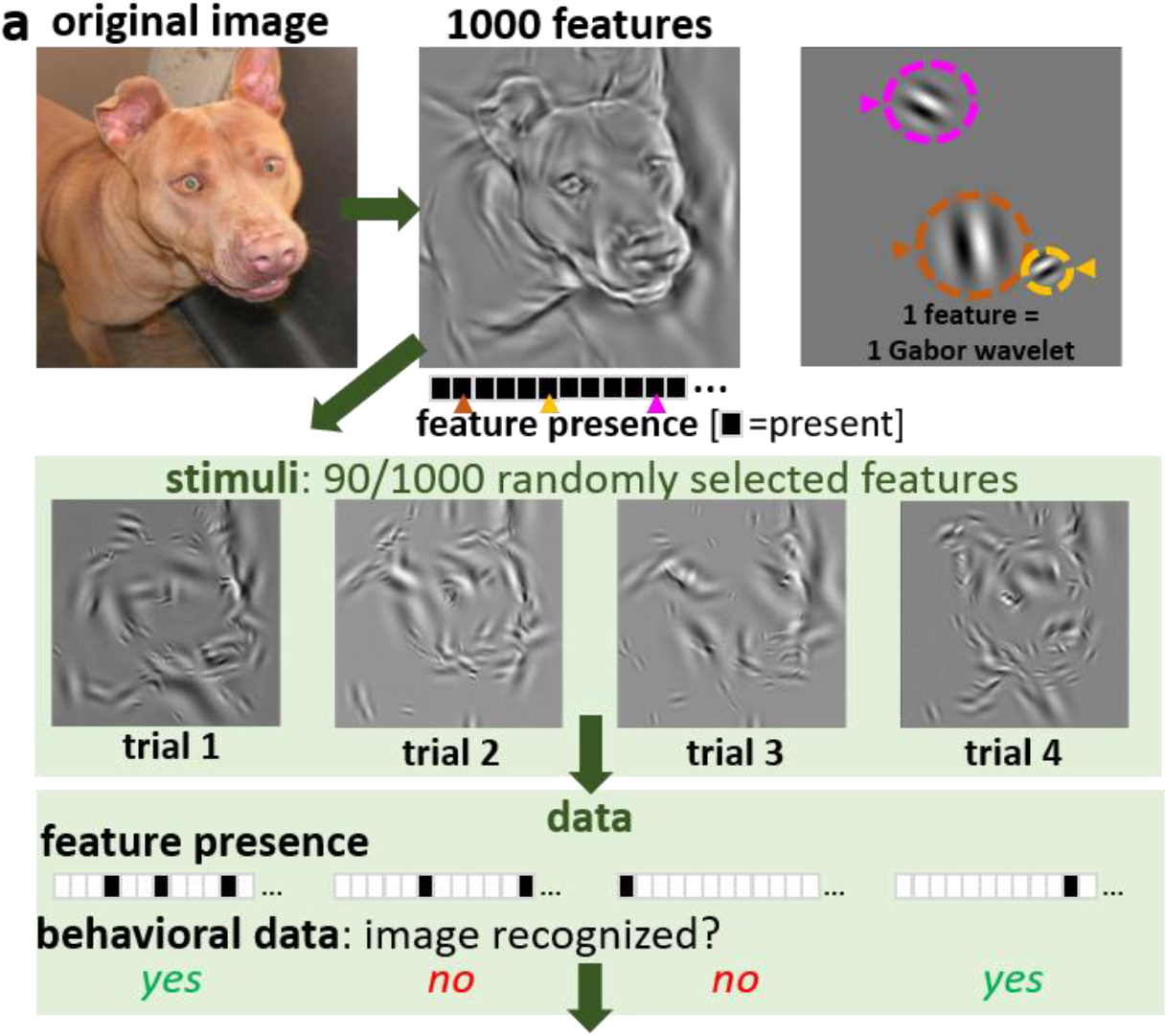
Feature diagnosticity mapping. a) Stimuli were reconstructed using the 1000 Gabor wavelets that best explained the pixel intensity in the original images. A random subset (90) of these features was presented during each trial while participants judged whether the image depicted a cat or a dog. A feature diagnosticity index (FDi) was computed based on the behavioral responses using an approach similar to reverse correlation. b) Images depicting the sum of the 200 features with highest (best features) and lowest (worst features) FDi values. c) graph showing that FDi value patterns replicate between participants (see methods for more details).

On average, participants recognized 49.7% (SD = 15.6%) of the partial reconstructions. A repeated-measure ANOVA revealed that there was no difference in recognition performance between cat and dog images (50.9% and 48.4% resp., F(1,100) = 0.50, p = .48), no effect of AQ group on recognition performance (high AQ group: 47.4%; low AQ group: 51.9%; F(1, 100) = 2.49, p = .12), nor an interaction between these two factors (F(1, 100) = 1.23, p = .27).

To quantify the relative importance of visual features we computed a feature diagnosticity index (FDi) for all 10,000 features – based on the average recognition accuracy for trials containing each feature. To reduce inter-participant and inter-image effects, averages were computed after z-scoring recognition accuracies within each participant and image – across the 1000 features of each image. If FDi values truly reflect feature importance for image recognition, images resulting from the summation of features with the highest FDi values should be most recognizable. Indeed, images reconstructed from the 200 features with the highest FDi values were easier to recognize than images reconstructed from the 200 features with the lowest FDi values (as shown for one exemplary image in Figure 1c). The efficacy of our method was further validated by the fact that the pattern of FDi values across all 10,000 features replicated significantly between participants (Pearson rho = .081, p < .0001 – permutation-based test, Figure 1c).

After having established that FDi values reliably measure the importance of features for image recognition, we tested if FDi values are elevated for high-spatial frequency features in high-AQ individuals. To this end, we grouped features into five equally-sized bins with ascending feature spatial frequencies. We tested for an interaction between spatial frequency and AQ group using a repeated-measure ANOVA with average bin FDi values as the dependent variable. This analysis revealed a main effect of spatial frequency (F (4,250) = 3.97, p < .005, Figure 2), and an interaction between spatial frequency and AQ group (F (4,250) = 4.12, p < .005, Figure 2). The interaction was mostly driven by the high-AQ group having elevated FDi values for the highest spatial frequency bin (.005 and -.013 resp.; t(50) = 3.80,p < .001). Therefore, our results are consistent with high-AQ individuals relying more on local details for image recognition.

**Figure 2:**
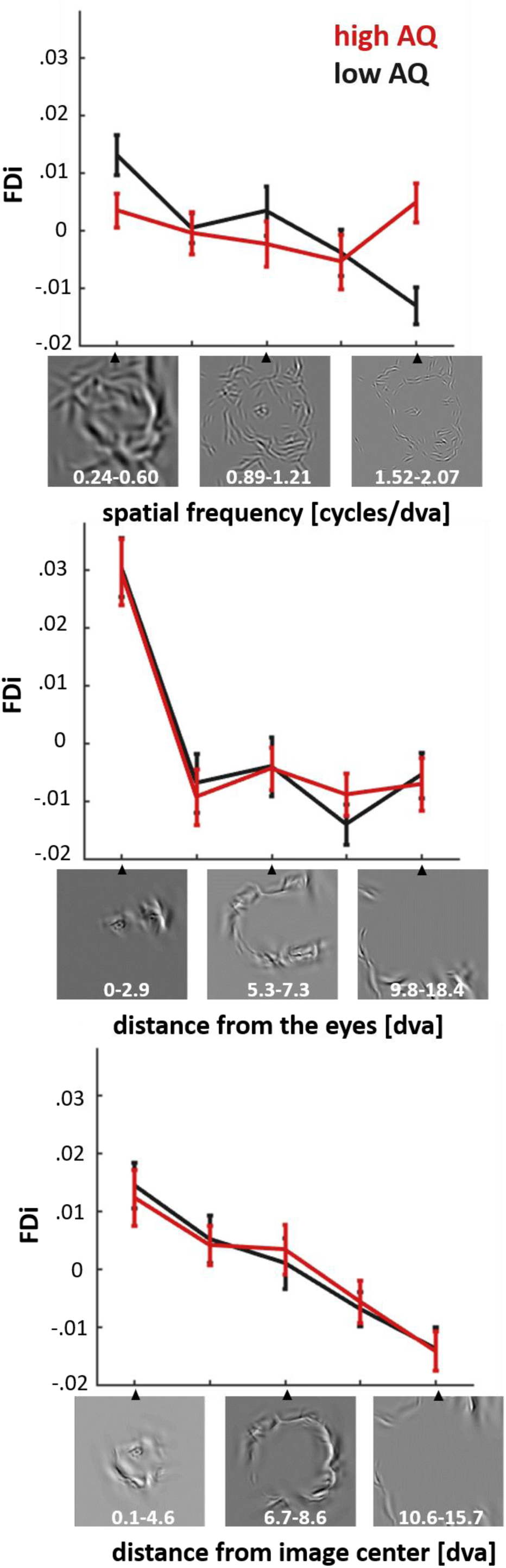
Effect of the number of autistic traits on the use of different types of visual features. The left panel depicts how feature diagnosticity is affected by AQ group (high AQ = AQ>14, low AQ = AQ<15)and feature spatial frequency. A clear interaction was observed between these two factors, which revealed that high-AQ individuals rely more on high spatial frequency information when recognizing cat and dog images. We also assessed the importance of the distance of features from the eyes (middle panel) and the image center (right panel) but found no interaction between these factors and AQ group.

Previous behavioral studies have linked ASD to reduced gaze durations for the eye-region in human faces^7–9^, and increased gaze durations for the central area of images^10^. To assess this, we repeated the analysis while grouping features into five ascending bins according to feature distance from the nearest eye or distance from the image center. This analysis revealed two main effects which indicate that feature diagnosticity decreases as a function of the distance from the nearest eye (F (4,250) = 27.4, p < .00001, Figure 2) and distance from the image center (F (4,250) = 14.3, p < .00001, Figure 2). Importantly, neither of these effects was modulated by AQ group. Therefore, our data does not replicate previously reported ASD effects on image center and eye-region processing. It is important to note that in contrast to these previous studies, our results are not based on eye movements nor on data from clinically diagnosed ASD patients.

Does our finding of an increased reliance-on detail for visual recognition in high AQ individuals generalize to individuals with an ASD diagnosis? Our study does not provide direct evidence for this as we measured a neurotypical student population. However, we are able to provide indirect evidence by testing if visual detail reliance depends most on the presence of clinically diagnostic AQ traits (with ‘trait’ we refer to a positive score on one of the 50 items of the AQ questionnaire). To this end, we quantified the clinical diagnosticity the 50 autism traits as the (natural) log of the trait prevalence ratio between clinically diagnosed ASD individuals and neurotypical students – based on previously published prevalence data^6^. E.g. the highest log odds ratio (2.04) was assigned to the trait measured with the item “*I enjoy social occasions*”, which ASD individuals disagree with 7.7 times more often than neurotypical students^6^. In addition, we quantified the reliance-on-detail for each participant as the linear regression coefficient between their average FDi values for each spatial frequency bin and the ascending bin numbers (1 to 5, see Figure 2). Subsequently, we performed a robust regression analysis that confirmed our hypothesis by revealing that our reliance-on-detail measure was best predicted by the presence of the most clinical diagnostic autistic traits (slope = .0023, t(48) = 3.40, p < .005, Figure 3).

**Figure 3:**
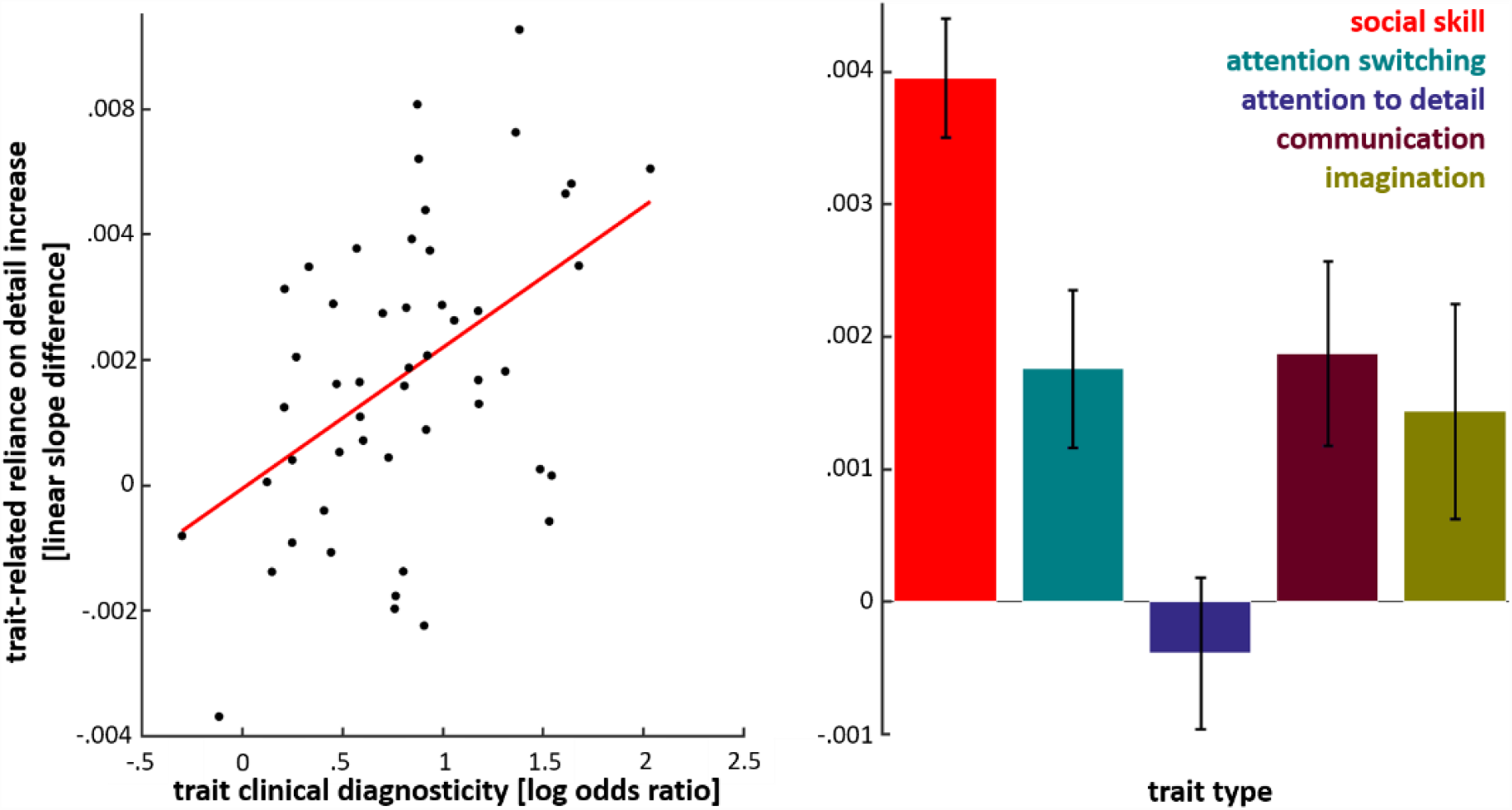
Reliance-on-detail effects scale with the clinical relevance of autistic traits. We assessed how a positive score on each of the 50 AQ items (referred to as ‘traits’) increased our participants preference for high spatial frequency features. Reliance-on-detail was quantified for each participant as the linear regression coefficient between their average FDi values for each of the spatial frequency bins and the ascending bin numbers (1 to 5, see Figure 2). Clinical diagnosticity of traits were quantified as the (natural) log of the trait prevalence ratio between clinically diagnosed ASD individuals and neurotypical students. The scatter plot shows that the clinical diagnosticity of traits scales positively with how much the presence of traits increases reliance-on-detail (left) Furthermore, reliance-on-detail increases were found to depend on the trait type (right), with the presence of “social skill” traits leading to above-average increases and “attention to details” traits to below-average increases.

In addition, we tested if trait-related increases of reliance-on-detail depend on the five different trait types (social skill, attention switching, attention to detail, communication and imagination). A one-way ANOVA revealed an effect of trait type (F(4,45) = 5.89, p < 0.001). Five post-hoc t-tests (bonf. corrected) testing for a difference between each trait type versus all others revealed that the presence of ‘social skill’ traits increase reliance-on-detail more than all other trait types (t(48) = 3.72, p < .005) while, surprisingly, ‘attention to detail’ traits give rise to a below-average effect (t(48) = –3.48, p < .01).

Finally, we analyzed reaction times (M = 895 ms, SD = 313 ms). We found that high-AQ participants took longer to respond than low-AQ participants (993 ms vs 804 ms, t(50) = 2.25, p < .05). Unfortunately, reaction times were found to carry no information about the relative importance of visual features for image recognition as reaction time based FDi values did not replicate across participants (Pearson rho = .0077, p = .14, permutation-based test).

In summary, we have developed a method that reliably measures the relative contribution of low level visual features to image recognition. With this method, we obtained the first direct evidence for natural image recognition depending more on high spatial frequency features in high-AQ individuals. This suggests that natural object recognition in high-AQ individuals is driven more by the fine details of objects as compared to individuals with a lower AQ. This effect was found to be driven most by positive scores on AQ items with the highest clinical relevance. Therefore, our results are likely to generalize to diagnosed ASD individuals. Lastly, our study suggests that AQ driven reliance-on-detail depends most on autistic traits related to social skills.

We did not, however, observe reduced processing of the eye-region, nor an increased processing of the central area of images – two effects previously associated with ASD^7,8,10^. Future studies are needed to resolve whether the absence of this effect is related the fact that we investigated ASD by comparing individuals with above- and below-median AQ scores. This would shed light on whether previously reported eye-region and image-center biases are specific to individuals that have been clinically diagnosed with ASD. Another intriguing possible explanation for this discrepancy could be that ASD leads to elevated gaze durations to the image center but that this does translate into object recognition relying more on the central image features. This insight could be obtained by combining our novel psychophysical paradigm with eye-tracking.

In conclusion, our results imply that the number of autistic traits we have increases the extent to which our day-to-day vision is driven by details. As this effect increases with the clinical relevance of the autistic traits, it appears likely that this effect generalizes to individuals with a clinical ASD diagnosis. Therefore, this study raises the possibility that a wide range of real-life abilities and difficulties associated with ASD relates to an increased reliance on visual details.

## METHODS

### Participants

52 healthy student volunteers (11 male, 41 female: average age = 19.4, SD = 1.05) with normal or corrected-to-normal vision took part in this experiment. All participants gave their informed consent after being introduced to the experimental procedure in accordance with the Declaration of Helsinki. The experimental procedure was approved by the ethics committee of the University of Birmingham (ethics reference ERN_15-1374P).

### Stimuli

Portrait images of five cats and five dogs were converted to 250×250 grey-scale images. We then used a custom-made algorithm, implemented in Matlab 2016a, to reduce images to 1000 Gabor wavelets. The aim of this algorithm was to find a set of wavelets which sum is able to describe most of the coarse and fine image details (see Figure 1a for a visualization of the features selected for one of the images). Wavelets considered had 29 (*n* = [1, 2, …, 29]) exponentially increasing spatial frequencies (*sf*) between 0.24 and 2.07 cycles/visual degree angle: 

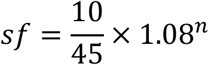

Wavelets were considered with 18 equidistant orientations between 0° and 180° (0° to 170° in steps of 10°). Features were selected iteratively from the lowest to the highest spatial frequency. During the first iteration, wavelets were selected based on the original grey-scale image. For consecutive iterations, the input image was a residual image resulting from least-square regression of the input to the previous iteration while using previously selected wavelets as regressors. During each iteration, eighteen Gabor wavelet filters (one per orientation) were applied to the input image. The output of this analysis enabled us to find the optimally-fitting orientation and phase for Gabor wavelets centered on each pixel of the input image for the current iterations’ spatial frequency. From these 250^2 wavelets, we considered only 25% with the highest amplitude. From these wavelets, we then randomly selected a number of wavelets (*nw*) increasing as a function of spatial frequency: 

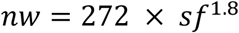

This resulted in 21 features being selected for 0.24 cycles/visual degree angle wavelets and 1008 features for 2.07 cycles/visual degree angle wavelets. During each iteration, we computed the covariance of each feature with the input image and discarded wavelets with covariances smaller than a fifth of the maximum observed covariance value. Finally, we selected 1000 wavelets from all spatial frequencies (from all iterations) having the highest covariance with the original grey-scale image. Amplitudes where set to an equal value for all wavelets. Partial reconstructions of the images were created by randomly selecting 90 wavelets from the set of 1000 and summating them (see Figure 1a). The pixel intensity range of the resulting images was kept constant, covering the full 0-255 range.

### Experimental procedure

Participants viewed partial reconstructions on an LCD from 70cm distance (visual degree angle (°), horizontal and vertical dimensions of the screen: 51.6° × 30.4°). Each trial started with a grey screen and a central fixation cross which participants were instructed to fixate, which we presented for a duration between 250 ms and 400 ms. Afterwards, a partial reconstruction was presented on a central area of the screen - covering 22.5° × 22.5° - and remained on the screen until a button press was made. Participants used their right hand to press one of the three available buttons with which they indicated having recognized a cat, a dog or that they weren’t sure about the type of animal they were shown. The next trial started as soon as a button was pressed. Due to the self-paced nature of the paradigm, participants completed a variable number of sessions - each consisting X trials per image - during the experimental session (6, 5, 4, 3 and 2 sessions were completed by 2, 14, 25, 10 and 1 participants resp.). Stimuli were presented and behavioral data was recorded using Matlab 2016a and the PsychToolbox (version 3).

### Data analysis

First, we assessed for each trial whether the image depicted was recognized. Then, we computed the average recognition performance for each participant and image feature by computing the average performance for trials containing the respective feature. This provided us with a three-dimensional matrix of recognition performances with the dimension number of participants (52), number of images (10) and number of features (1000). Next, we obtained our feature diagnosticity index (FDi) values by z-transforming the performance values within each participant and image. This final step is important because it precludes FDi values being higher for features of images that are easier to recognize. Furthermore, it ensures that participants’ relative contributions to the following analyses do not depend on their average recognition performance nor the variability of their responses.

To evaluate the replicability of the observed FDi values, we randomly split the data into two halves (two times 26 participants) 100 times and computed the Pearson correlation between the average FDi values across splits. Replicability was then measured as the average Pearson correlation value across these 100 splits. The probability of observing this value by chance was determined by computing a null-distribution by re-computing this value 10,000 times while permuting the relative feature labels across splits.

To assess effects of spatial frequency, distance from the nearest eye and distance from image center on FDi values, we created five equally sized ascending bins based on each of these parameters. Thereafter, we computed the average FDi value within each of these bins separately for each participant. In addition, we assigned each participant to the high AQ group and low AQ group depending on whether their AQ was or was not higher than the median AQ across all participants. This enabled us to perform three 5 × 2 repeated measure ANOVAs with average within-bin FDi values as the dependent variable. Each ANOVAs second factor was AQ group while the first factor was the binning feature: spatial frequency, distance from the nearest eye or distance from image center.

